# Closed-loop spinal cord stimulation is superior in restoring locomotion in rodent models of Parkinson’s Disease

**DOI:** 10.1101/2022.09.19.508432

**Authors:** Benjamin Rees, Eleonora Borda, Miguel A. L. Nicolelis, Amol P. Yadav

## Abstract

Dorsal column stimulation (DCS) of the spinal cord is emerging as a promising new technology to treat Parkinson’s disease (PD). However, optimal stimulation settings that maximize its therapeutic effect on PD symptoms are yet to be determined. Here we demonstrate a closed-loop DCS (CLDCS) paradigm – a substantial advancement from previously tested continuous high-frequency DCS – in a bilateral intrastriatal 6-hydroxydopamine (6-OHDA) rodent model of PD. Firstly, CLDCS, triggered by corticostriatal beta frequency oscillations facilitated a pro-locomotion brain state that restored locomotion and reduced akinesia. Secondly, CLDCS was better at disrupting ongoing beta oscillations and achieved it with lesser overall charge delivery than continuous open-loop stimulation. These results indicate that CLDCS is markedly better than traditional spinal cord stimulation methods and can potentially be highly effective in treating PD symptoms. We envision that the CLDCS approach can be beneficial in the treatment of other neurological disorders which showcase similar pathological neuronal oscillations.

## Introduction

Parkinson’s disease (PD) is a movement disorder that causes progressive worsening of motor function leading to symptoms of tremor, rigidity, bradykinesia, and gait instability (1). Non-pharmacological methods such as deep brain stimulation (DBS) of the subthalamic nucleus (STN) and globus pallidus interna (GPi) improve symptoms of tremor, rigidity, and bradykinesia (2–4). However, DBS is not very effective in treating gait and balance abnormalities, leading to life-threatening complications such as ‘freezing of gait’ (FOG) and ‘falls’ (5, 6). Hence, other locations along the sensorimotor pathway such as the pedunculopontine nucleus (PPN) (7–9), the mesencephalic locomotor region (MLR) (10), and the spinal cord dorsal columns (11) have been proposed as alternative stimulation targets to treat the axial symptoms related to PD.

Epidural stimulation of the dorsal columns of the spinal cord – i.e., dorsal column stimulation (DCS) – was shown to be effective in improving motor symptoms in rodent and non-human primate models of PD (12–16). Subsequently, several case reports have also demonstrated that DCS improves gait abnormalities, especially freezing of gait (FOG) in advanced PD patients (17–20) – for a detailed list of studies, see (11, 21). Yet, the exact mechanism by which DCS alleviates symptoms is not fully understood. In addition, optimal DCS parameters which maximize therapeutic benefit in patients are yet to be determined.

Previous work showed that continuous high-frequency DCS (>300 Hz) was most effective in desynchronizing beta-oscillations – a well-established neurophysiological biomarker of PD (12, 14). However, from a clinical perspective, therapies involving continuous high-frequency stimulation can significantly drain the battery power of the implantable pulse generator, requiring frequent replacement surgeries (22). Hence treatments that maximize therapeutic benefit while delivering the least amount of charge to the neural tissue are ideally suited for clinical translation. Recently, adaptive DBS studies have shown considerable promise by incorporating real-time modulation of the stimulation amplitude or duty-cycle by a reliable neurophysiological biomarker such as gamma or beta band oscillations (23, 24). Based on these recent trends and with the goal of minimizing the charge delivered to the spinal cord, we hypothesized that closed-loop dorsal column stimulation (CLDCS) would be a superior therapeutic approach to treat PD symptoms.

To test our hypothesis, we developed a CLDCS paradigm that detects ongoing beta frequency oscillatory activity in the cortico-striatal circuits and delivers a train of DCS pulses to disrupt the ongoing oscillations (Figs 1a, 1b, 1c, 1d). We used a bilateral intra-striatal 6-hydroxydopamine (6-OHDA) lesion rat model of PD in our investigation. We hypothesized that closed-loop DCS would be better than, or at the very least comparable to, open-loop DCS (OLDCS) in ameliorating PD symptoms and desynchronizing beta-frequency oscillations. We expected that our novel stimulation paradigm would reduce the total charge delivered to the nervous system, thus potentially improving battery life when used in a clinical setting to treat PD patients.

**Figure 1:**
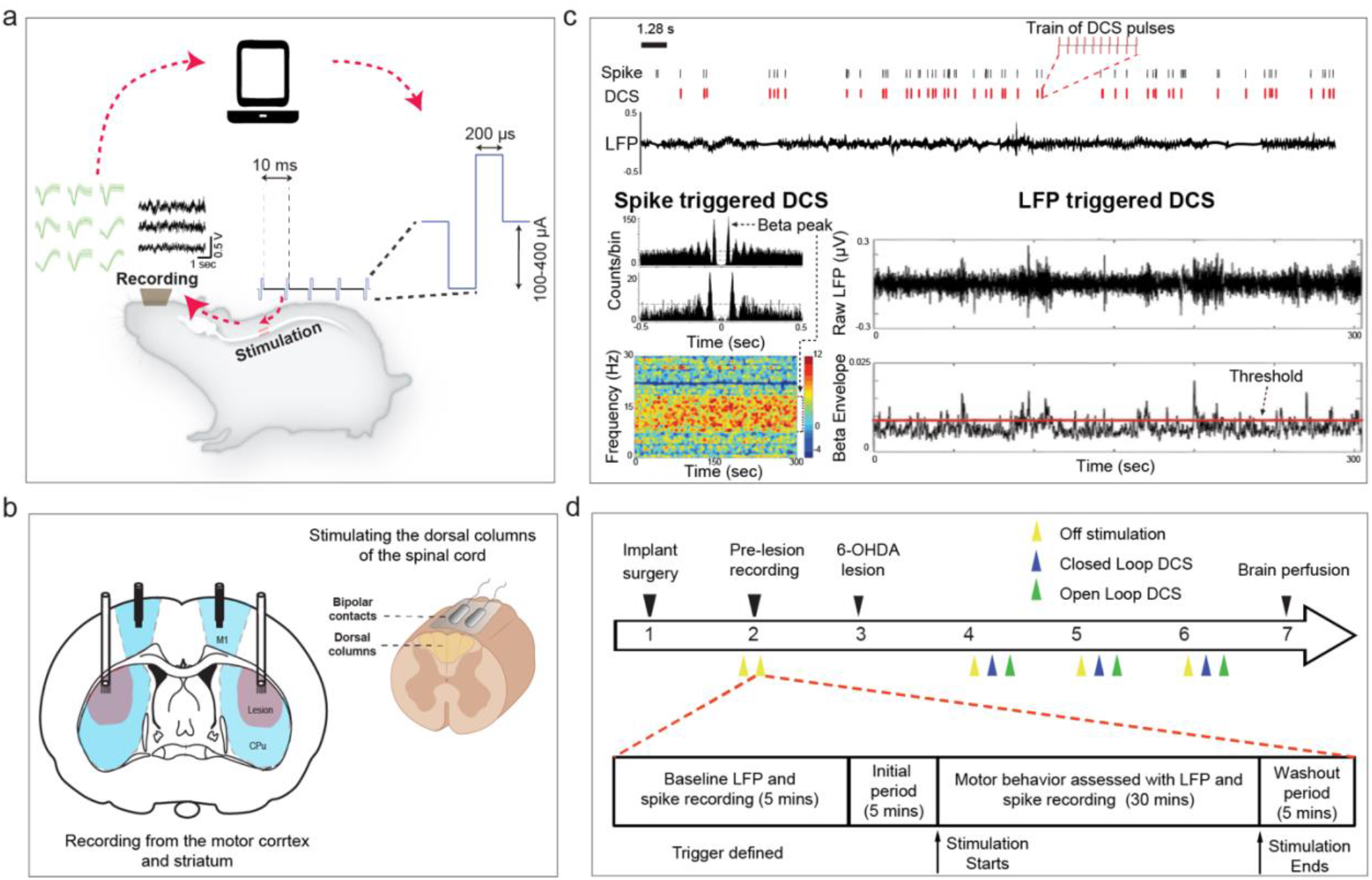
Closed-loop dorsal column stimulation (CLDCS). a,b) Schematic for CLDCS. Rats were implanted with stimulation electrodes on the epidural dorsal surface of the spinal cord and with recording electrodes in the primary motor cortex and dorsolateral striatum. Real-time processing of brain activity was performed to detect neurophysiological biomarkers, which were used to trigger stimulation (train of ten biphasic pulses at 100 Hz, 200 µs per phase). c) CLDCS was performed using two approaches: 1) spike-triggered – where striatal neurons with beta-frequency oscillatory activity were used to trigger stimulation; and 2) local field potential (LFP)-triggered – where stimulation was delivered whenever beta-frequency oscillations in the striatal LFPs crossed a preset threshold. d) One week after implant surgery, 6-hydroxydopamine (6-OHDA) injections were performed. One week after, experimental sessions began, comprising 5 minutes of baseline recording to identify spike or LFP channels for stimulation control. During 30 mins of assessment period (recording of motor and neuronal activity), either closed-loop DCS (CLDCS) or no stimulation (Off stim) was delivered. To compare the efficacy of CLDCS and open-loop DCS (OLDCS), in some sessions, the assessment period was increased to 45 mins, where CLDCS and OLDCS were delivered during two non-consecutive 5-min epochs in a randomized order preceded and succeeded by 5-min Off stim periods.

## Results

Rats were implanted with stimulation electrodes in the dorsal epidural space of the T3 spinal vertebra, recording electrodes in the motor cortex (M1) and dorsolateral striatum (Str) and guide cannulas in the dorsolateral striatum (Fig 1b). One week after implant surgery, 6-hydroxydopamine was infused in the striatum to induce Parkinsonian symptoms (Fig 1d). One week after lesioning, closed-loop dorsal column stimulation (CLDCS) experiments were initiated with simultaneous recording of neural activity (spike and LFP) and locomotion in the open field. CLDCS was delivered in two modes (Fig 1c): spike-triggered DCS (spiking of an M1 or Str neuron with beta oscillatory activity triggered a train of DCS pulses); and LFP-triggered DCS (beta oscillations in Str LFPs triggered a DCS pulse train).

### Closed-loop DCS improves locomotion and reduces beta-frequency oscillations in corticostriatal circuits

Bilateral injection of 6-OHDA in rats induced severe parkinsonism, demonstrated by symptoms of akinesia (Figs 2a, 2b), thereby reducing overall locomotion [distance traveled decreased 87% from pre-lesion and time spent in rest increased 76% from pre-lesion. p<0.001, two-sample t-test, Figs 2g, 2h], along with sustained increase in corticostriatal LFP beta-frequency oscillations (max increase 98% in right striatum, p<0.01, two-sample t-test, Figs 2d, 2e, 2k-n). CLDCS dramatically improved locomotion [distance traveled increased 204% from off-stim, p<0.001, and time spent in rest decreased 17% from off-stim, p<0.01, two-sample t-test, Figs 2c, 2g, 2h]. DCS-induced improvement in locomotion was accompanied by reduced corticostriatal LFP beta-frequency oscillations in the CLDCS condition compared to the off-stim condition (max decrease 61% in right striatum, p<0.001, two-sample t-test, Figs 2f, 2k-n). This indicated that the application of DCS in a closed-loop manner - triggered by underlying pathological neuronal activity associated with PD - led to desynchronization of corticostriatal low-frequency oscillations and thus alleviation of symptoms of akinesia.

**Figure 2:**
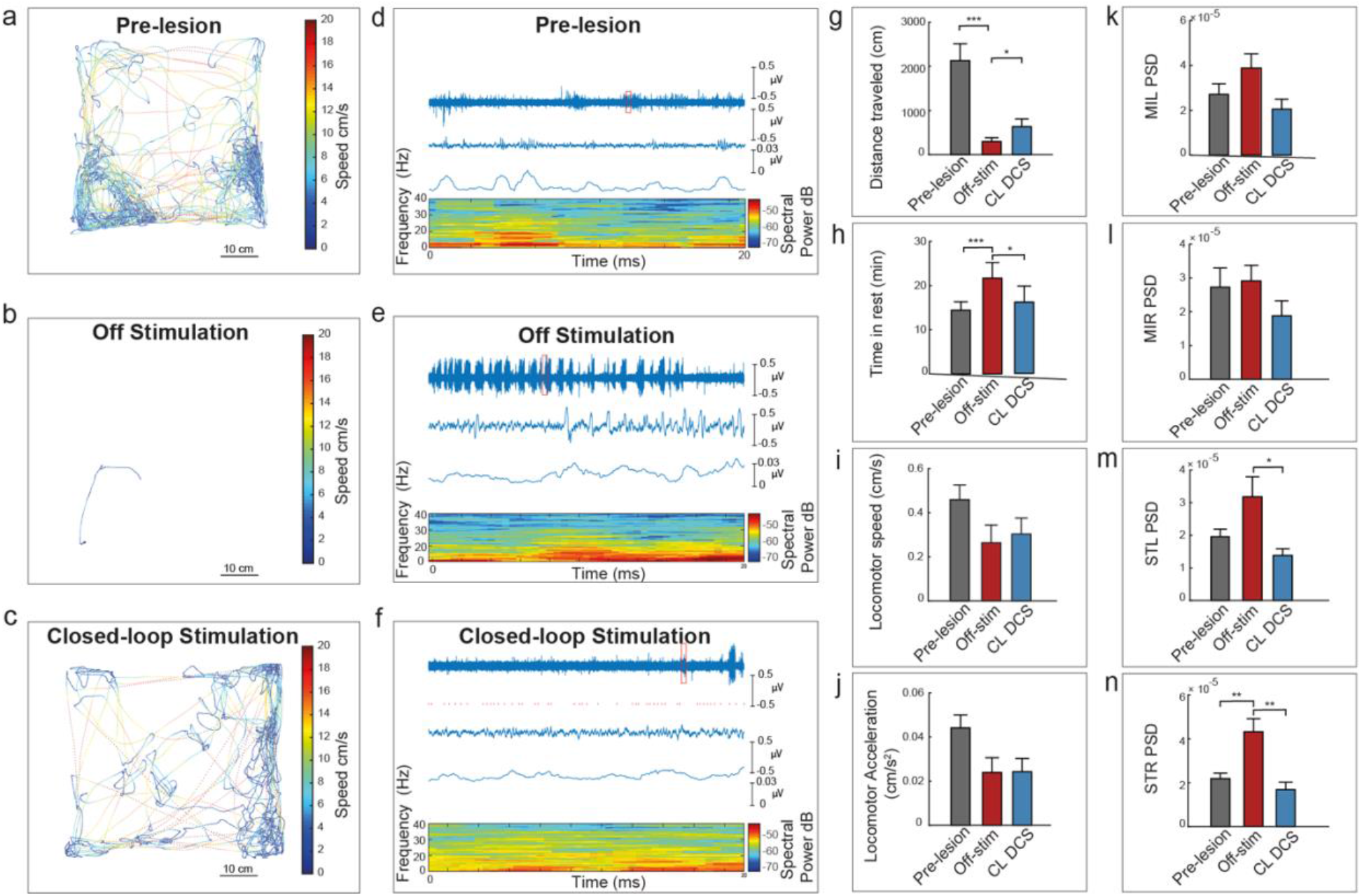
CLDCS improves locomotion and desynchronizes corticostriatal beta-frequency oscillations. a,b,c) Representative locomotion tracking of a rat in a pre-lesion (6-OHDA injection) session and post-lesion Off-stimulation and CLDCS session shows marked differences in spontaneous locomotor activity. Representative striatal LFP activity in pre-lesion session and post-lesion Off-stimulation and CLDCS sessions. Top trace shows raw LFP for entire session, second row shows zoomed LFP for 20-second window, third row shows filtered and rectified beta frequency, fourth row shows spectrogram. There were clear differences in the beta-frequency oscillations in the three representative sessions of a rat. g-n) Averaged locomotor activity (distance traveled and time spent in rest) and neural activity (beta-frequency oscillations in left (STL) and right striatum (STR) showed dramatic improvement in CLDCS sessions as compared to off-stimulation sessions across all animals (n=5 rats). Values represent mean ± sem, *:p<0.05; **:p<0.01; ***:p<0.001 (two-sample t-test)

### Closed-loop DCS creates a pro-kinetic brain state

To understand whether CLDCS influenced the firing rates of neurons, we looked at spiking activity across all neurons in six rats between off-stimulation and CLDCS conditions. We found that CLDCS significantly modulated firing rates. In 135 (23%) neurons, CLDCS increased firing rates, while in 216 (36.8%) neurons, CLDCS decreased firing rates significantly (Fig 3a). In neurons that showed changes in firing rates in the CLDCS condition, overall firing rates were significantly different (10.87 Hz vs. 8.91 Hz, neurons showing inhibitory effect, Fig 3b left; 6.16 Hz vs. 8.04 Hz, neurons showing excitatory effect, Fig 3b right; Wilcoxon matched-pairs signed rank two-tailed test) as compared to the off-stimulation state. While it was clear that CLDCS modulated overall firing rates across the session, to understand how it facilitated locomotion, we studied the population spiking activity of all neurons around locomotion initiation. We observed mixed neuronal responses to the onset of locomotion, where some neurons exhibited an increase in firing rate around locomotion initiation while others exhibited a decrease (Fig 3c). Principal component analysis of all neurons across all rats was performed to study the transition of the 3D neural trajectory before and during locomotion. Between off-stim and CLDCS conditions, clear differences were observed in the variance captured of the 3D neural states (measured as volume of fitted ellipsoid) and the total path length of the neural trajectory (Fig 3d). Overall, variance captured and path length were significantly smaller in CLDCS as compared to off-stimulation conditions (Fig 3e, Wilcoxon matched-pairs signed rank two-tailed test), suggesting that the population activity had an easier and more controlled transition from the pre-locomotion to the locomotion state during CLDCS.

**Figure 3:**
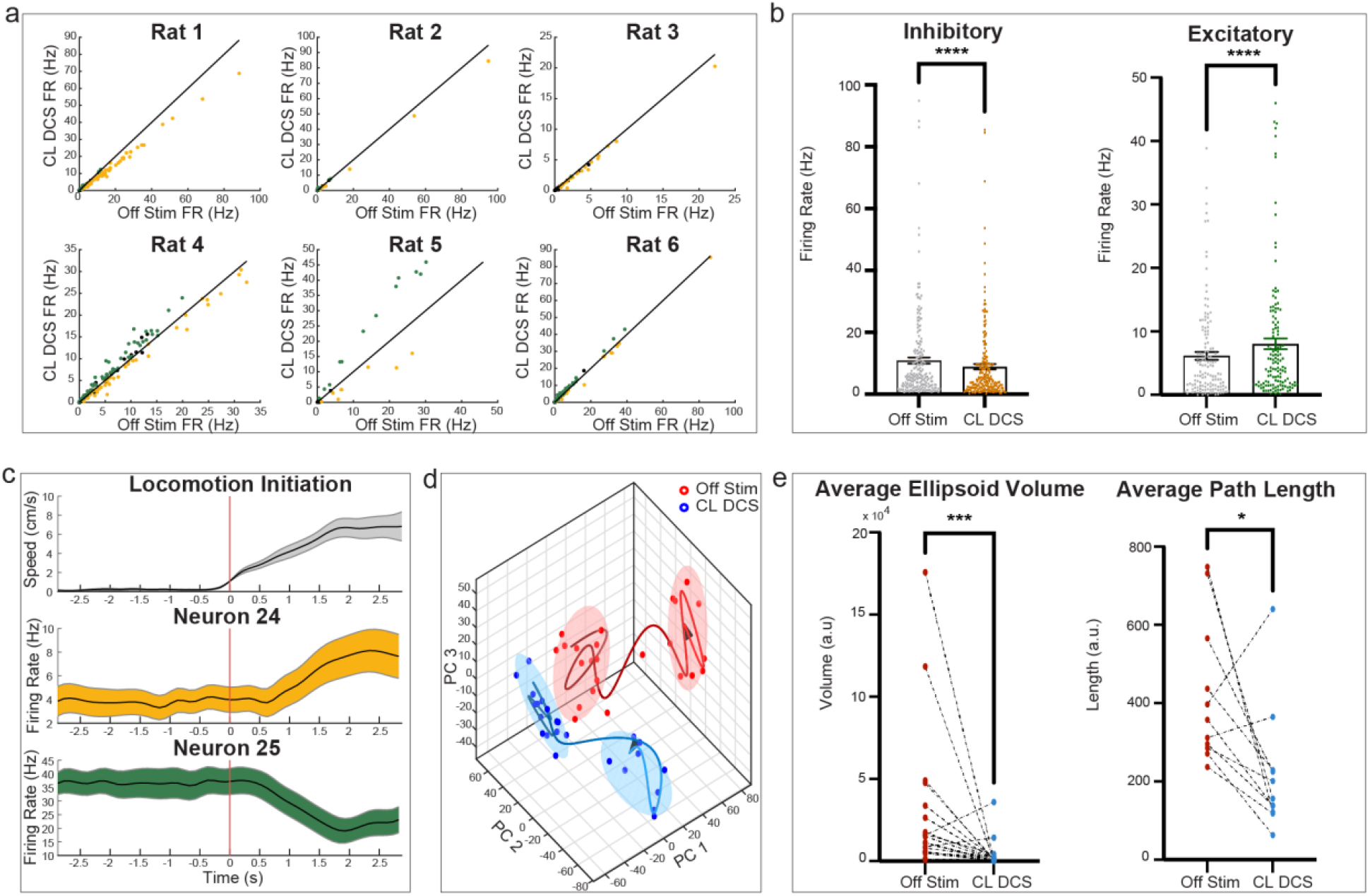
CLDCS creates a pro-kinetic brain state in Parkinsonian rats. a) CLDCS had a significant effect on firing rates (FR) of neurons in all rats as compared to baseline Off-stim period. Neurons showing significantly-increased FR (green, excitatory response), -decreased FR (yellow, inhibitory response), and no significant FR change (black) are shown for each rat. b) Among all neurons that showed either significant excitatory or inhibitory response to CLDCS, the overall firing rate for the entire session was significantly different between CLDCS and Off-stim periods (Wilcoxon matched-pairs signed rank test). c) Representative neurons showing increase (neuron 24) and decrease (neuron 25) in FR around locomotion initiation. d) Neural trajectory of first three principal components after principal component analysis of firing rates of all neurons (±3 seconds around locomotion initiation). Ellipses fitted to pre-locomotion and during-locomotion trajectory points show disjoint clusters of different sizes and path lengths between Off-stim (orange) and CLDCS (blue) conditions. Fitted ellipse volume and path length of neural trajectory were significantly smaller between Off-stim and CLDCS across multiple sessions (n=11, Wilcoxon matched-pairs signed rank test), suggesting that the transition from pre-locomotion to locomotion state was easier during the CLDCS condition as compared to Off-stim.

### Closed-loop DCS is superior to open-loop DCS

To determine whether closed-loop stimulation was better than open-loop stimulation, we conducted experiments where both CLDCS and OLDCS paradigms were delivered within the same session (n=15 sessions) in a random order so that their effect on locomotion and neuronal activity could be directly compared. Our results demonstrated that CLDCS was significantly better at increasing distance traveled and decreasing time spent in rest as compared to OLDCS [343 ± 45 cm vs. 244 ± 41 cm (p<0.001); 237 ± 19 sec vs. 341 ± 32 sec (p<0.05); one-way ANOVA followed by Tukey’s multiple comparisons test, Figs 4a, 4b]. In parallel, CLDCS reduced cortical and striatal LFP beta oscillations significantly better than OLDCS at all recording sites (max 53% better in left striatum, one-way ANOVA followed by Tukey’s multiple comparisons test, Figs 4e-h). Thus, our results indicated that CLDCS was superior to OLDCS in desynchronizing beta frequency oscillations, thereby promoting locomotion in Parkinsonian rodents.

**Figure 4:**
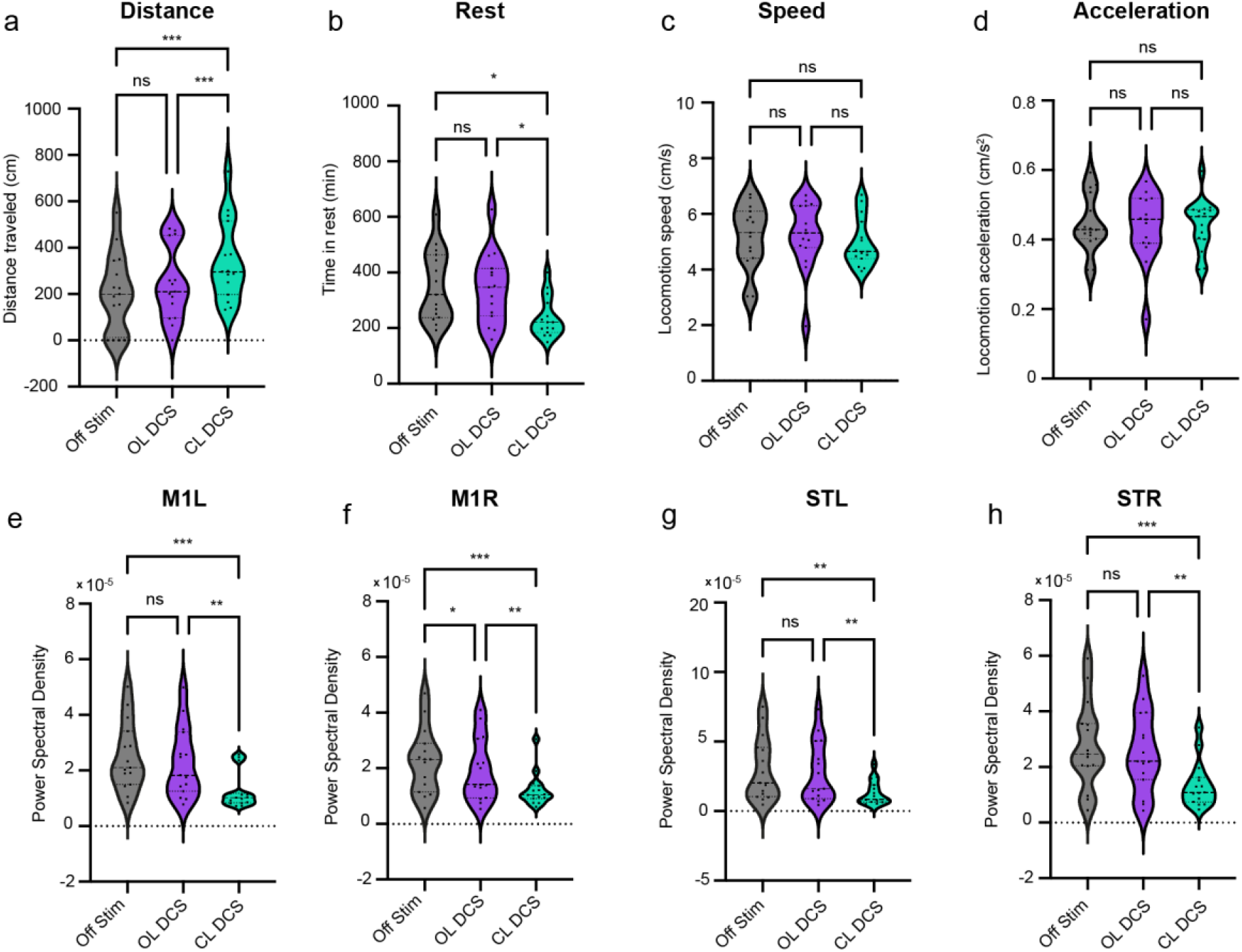
Closed loop DCS is superior to open-loop DCS. CLDCS was significantly better than OLDCS in improving locomotion [distance traveled (a) and time spent in rest (b)] and reducing pathological beta frequency oscillations in motor cortex (e,f) and striatum (g,h). There was no significant difference in locomotion speed and acceleration across all conditions (c,d). Violin plot shows median and quartiles. *: p<0.05; **: p<0.01; ***:p<0.001 (One-way ANOVA followed by Tukey’s Multiple Comparisons test)

### Final PCA of all data

Finally, to determine how the measured parameters (locomotion and LFP beta power) were influenced by various paradigms (pre-lesion off-stim, OLDCS, and CLDCS), we took the principal component scores calculated from all sessions and all rats and measured the Euclidean distance from the pre-lesion cluster (Figs 5a, 5b). The CLDCS cluster was significantly closer to the pre-lesion cluster as compared to OLDCS (2.82 ± 0.26, p<0.01) and off-stim (3.04 ± 0.28, p<0.05) clusters (one-way ANOVA followed by Tukey’s multiple comparisons test, Fig 5c)). This indicated that although CLDCS didn’t completely restore the rat’s behavioral and neural state to how it was before developing Parkinsonian symptoms, it did a better job than OLDCS and off-stim. Overall, CLDCS achieved higher efficacy in improving symptoms with lesser total charge delivered. The average stimulation frequency across all animals and all sessions was 18.3 ± 1.84 Hz compared to 100 Hz for OLDCS, suggesting a >80% reduction in the total charge (Fig 5d).

**Figure 5:**
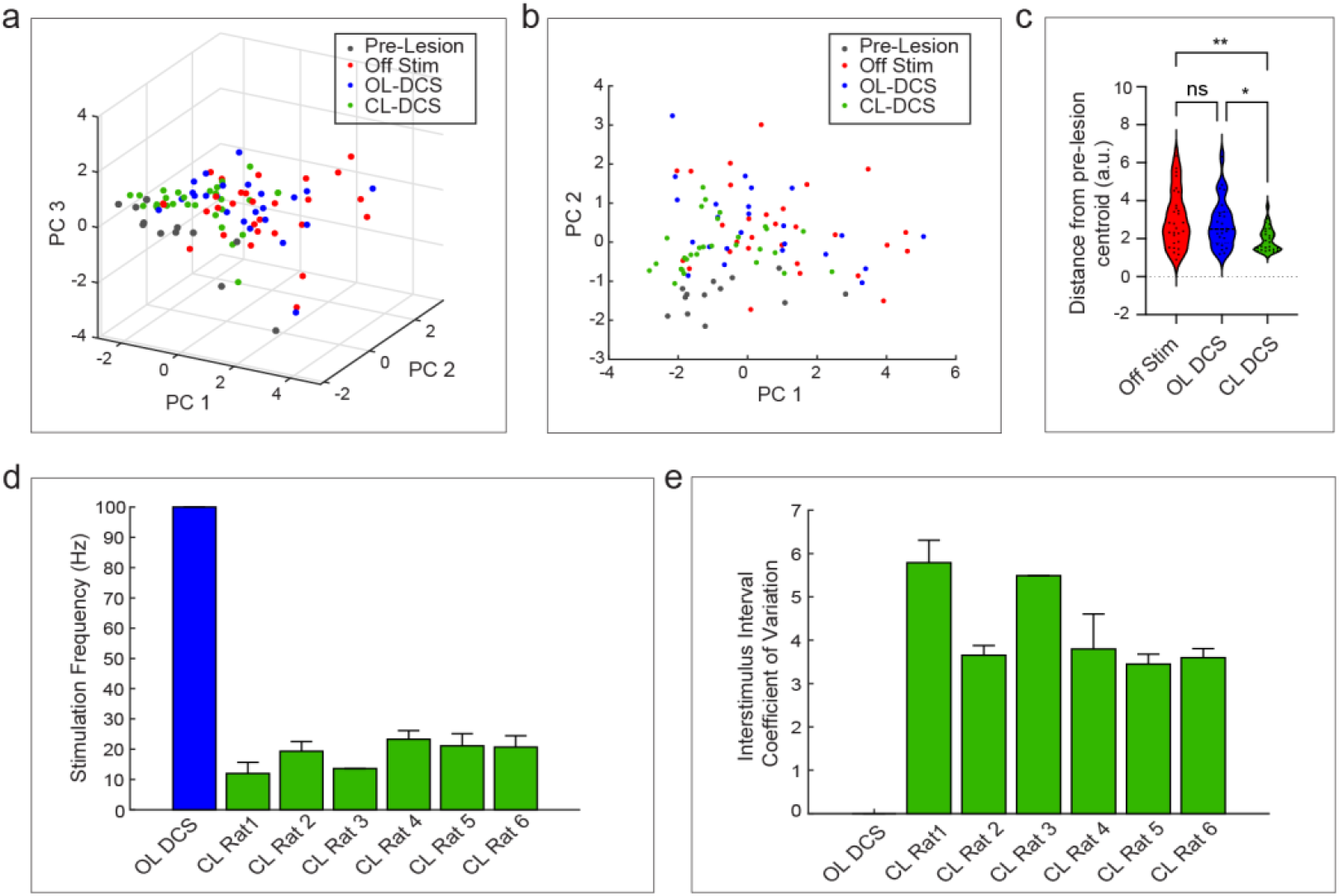
Closed loop DCS brings the animal’s brain and behavior closest to pre-lesion state as compared to open loop DCS and off-stimulation. Principal component analysis of locomotion and beta frequency power parameters [First 3 PCs (a) and first 2 PCs (b) revealed that points representing CLDCS condition (green) clustered closer to pre-lesion points (black) as compared to OLDCS (blue) and Off-stim (red). c) Violin plots showing distance from pre-lesion centroid (median and quartiles) for all points in (a) indicate that CLDCS points were significantly closer to the pre-lesion cluster than OLDCS and Off-stim. d,e) Average stimulation frequency and average coefficient of variation of interstimulus interval across all sessions in each rat for OLDCS (blue) and CLDCS (green) conditions show that CLDCS creates highly random stimulus patterns at a low average frequency as compared to OLDCS, thereby reducing total charge delivered.

## Discussion

We demonstrated that closed-loop stimulation of the dorsal columns of the spinal cord (CLDCS) triggered by beta oscillations in the corticostriatal neural activity improved spontaneous locomotion and, in turn, reduced overall corticostriatal LFP oscillations in a bilateral 6-OHDA rat model of PD. CLDCS was superior to open-loop DCS (OLDCS) in improving symptoms and desynchronizing corticostriatal brain activity. Although CLDCS was more effective in ameliorating symptoms, it achieved this feat using a much lower average stimulation frequency than OLDCS. Our results showcase a critical step in advancing DCS as an alternative therapeutic approach for treating Parkinsonism.

So far, previous pre-clinical and clinical studies exploring spinal cord stimulation for PD have used continuous stimulation. For instance, in rodents and marmoset monkeys, high-frequency stimulation at 300 Hz was found to be most effective in improving symptoms of locomotion and gait (12–14), while studies focusing on the neuroprotective effect of DCS in rats found continuous DCS at 50 Hz to be most effective. Similarly, several clinical studies have reported improvements in gait and posture symptoms in PD patients using continuous stimulation ranging from 60 Hz to 300 Hz [for a detailed review, see (11, 21)]. As such, the optimal DCS parameters that address PD symptoms are still to be determined.

The temporal pattern of a stimulus train is an important feature that has recently gained interest in DBS research (22, 25). However, initial studies using irregular patterns of DBS showed mixed results when compared to regular DBS in improving tremor and bradykinesia in PD patients (26–28). In contrast, adaptive DBS, where STN stimulation onset/termination was controlled by 3-35 Hz STN LFP magnitude or STN stimulation amplitude was modulated by 60-90 Hz motor cortex oscillations indicative of dyskinesias, showed remarkable superiority over conventional DBS along with significant reduction (38-56%) in stimulation time or energy consumption (23, 24, 29). These initial findings suggest that symptom amelioration can be enhanced by driving the brain circuits at a temporal rate that facilitates motor function or by modulating stimulation using pathological brain activity, which is correlated with symptom onset/intensity. Based on our primary hypothesis that DCS desynchronizes supraspinal brain circuits involved in pathological low-frequency oscillations by simultaneous activation of fibers in the dorsal column (DC) medial lemniscal pathway (11), here we can speculate that CLDCS might lead to activation of a higher number of DC fibers due to its highly aperiodic nature. This can explain why we observed greater low-frequency desynchronization during CLDCS compared to OLDCS. Indeed, in our case, the overall CLDCS stimulus train was highly random (interstimulus interval coefficient of variation was greater than three as compared to zero for OLDCS, Fig 5e). A relation between DCS-induced sensory perception thresholds and stimulation frequency was demonstrated earlier (30). In other experiments, we observed that sensory thresholds decrease as stimulus aperiodicity increases (data not shown here), suggesting that stimulus aperiodicity may directly influence the number of dorsal column fibers activated by DCS. In future studies, we plan to compare CLDCS with OLDCS patterns at different randomness levels to isolate and understand the role of stimulus periodicity in symptom alleviation.

We showed that CLDCS had both inhibitory and excitatory effects on the spiking activity of neurons in the motor cortex and striatum (Fig 3a, 3b). A similar effect of open loop stimulation has already been established in previous research (14, 31). In addition, by performing dimensionality reduction on the time-series population activity of all neurons before and during locomotion bouts, we could represent aggregate neural activity using principal components (PC) as locomotion unfolded over time. In both the Off-stim and CLDCS conditions, two neural activity states within the PC space were observed across bouts of locomotion marked by distinct clusters for a pre-locomotion and a locomotion state. Relative to CLDCS, the Off-stim condition showed greater 3D spread for both activity states, suggesting reduced variance in the neural trajectory over time because of stimulation (Figs 3d, 3e). Although this reduction is only correlative, it is possible that the dampening of variance during the time leading up to locomotion puts the brain in a more favorable state to initiate locomotion, indicated by the reduced path length of the neural trajectory (Figs 3d, 3e). Though the neural population dynamics underlying spontaneous locomotion are not heavily investigated, recent evidence highlights that hind-limb kinematics and gait phase during unconstrained locomotion are represented in the low-dimensional state space models of motor cortex population activity (32, 33). Future studies could identify low-dimensional latent variables that encode spontaneous locomotion initiation and study whether these variables can be used to modulate CLDCS to create a pro-locomotion brain state.

Our CLDCS protocol achieved a >80% reduction in overall stimulation frequency (Fig 5d). From a clinical translation perspective, reducing delivered stimulation energy has multiple benefits. For instance, it can decrease stimulation-associated side effects, prolong implantable pulse generator battery life, prevent daily charging, and allow miniaturization of rechargeable pulse generators. Even though the clinical implementation of DCS for PD treatment is in its infancy, we can envision a future where both DBS and DCS can be implemented in the same patient. While DBS can focus on treating tremors and bradykinesia, concurrently, DCS can be applied to treat axial symptoms affecting gait, balance, and posture. Indeed, implementing a closed-loop DCS protocol utilizing DBS electrodes for sensing would significantly advance the clinical translation of this novel technology. Furthermore, we foresee that our CLDCS paradigm could be applied to treat several neurological conditions such as epilepsy, stroke, brain injury, spinal cord injury, and other neuropsychiatric disorders which manifest robust neuronal biomarkers.

## Materials and Methods

All animal procedures were performed according to prior approved protocols by Duke University Institutional Animal Care and Use Committee and in accordance with the National Institute of Health Guide for the Care and Use of Laboratory Animals (NIH Publications No. 80–23).

### Surgery for microelectrode array and stimulation electrode implantation

Animals went through two different implantation procedures performed in the same surgery under general anesthesia: one to implant recording electrodes in the brain and the second to implant stimulation electrodes on the spinal cord (Fig 1b). Anesthesia was induced with isoflurane and maintained with ketamine (100 mg/kg), xylazine (10 mg/kg), and atropine (0.05 ml). Movable recording microelectrodes were implanted stereotactically bilaterally in the rats’ primary motor cortex (M1) and dorsolateral striatum (STR). Stimulation electrodes for spinal stimulation were implanted in the dorsal thoracic epidural spinal cord space under thoracic vertebra T2-T3 and tied to the spinous process with a surgical suture to prevent electrode migrations (13).

Postoperative weight was monitored daily. 6-OHDA hydrobromide (Sigma Company, USA - 3.5 mg/ml in 0.05% ascorbate saline) was injected bilaterally into the dorsolateral striatum using a needle through implanted cannulas driven by a syringe pump (Sage, Model 361, Firstenberg Machinery Co Inc., USA) via 10 µL Hamilton syringe, at 7 µL/min. The needle was left in situ for 5 minutes and withdrawn slowly to prevent backtracking of the drug. Destruction of noradrenergic fibers and terminals was prevented by intraperitoneal injection of 1,3-Dimethyl-2-imidazolidinone (DMI, Sigma Company, 25 mg/kg) 30 minutes before 6-OHDA treatment.

### Experimental design

One week after surgery, locomotion and brain recording were performed to determine pre-6-OHDA lesion baseline. Two weeks after surgery, 6-OHDA was infused bilaterally in the dorsolateral striatum to induce Parkinsonism. Once Parkinsonian symptoms were validated by lack of movement and abnormal posture and gait, as established previously (13), stimulation experiments were initiated (Fig 1d). A typical experimental session included five minutes of baseline recording followed by thirty minutes of assessment period. Spikes or LFP channels to be used for brain-triggered stimulation (train of ten biphasic pulses at 100 Hz, 200 µs per phase, 100-400 µA, Fig 1a) were identified from the baseline recording. In spike-triggered CLDCS, striatal neurons with beta-frequency oscillatory activity were used to trigger spinal stimulation, and in (LFP)-triggered CLDCS, stimulation was delivered whenever beta-frequency oscillations in the striatal LFPs crossed a preset threshold (Fig 1c). During the assessment period, either CLDCS or no stimulation (Off stim) was delivered along with recording of locomotion and neuronal activity. In some sessions, the assessment period was increased to forty-five minutes to compare the efficacy of CLDCS and open-loop DCS (OLDCS). During this period, CLDCS and OLDCS were delivered in two non-consecutive five-minute epochs in a randomized order preceded and succeeded by five-minute Off stim periods.

### Closed-loop control

To detect spike channels with beta oscillations autocorrelation and spectrograms were used to determine oscillatory peaks during the five-minute baseline recording (Fig 1c). Stimulation was delivered (train of ten pulses at 100Hz) every time the selected single unit spiked. To detect LFP channels with beta oscillations, raw LFPs from five-minute baseline recordings were filtered in the beta range (10-25 Hz), rectified, smoothed, and a threshold was set as mean plus standard deviation (Fig 1c). Stimulation was delivered (train of ten pulses at 100 Hz) every time selected LFP data crossed the set threshold. To prevent continuous stimulation, a refractory period of 100 milliseconds was inserted at the end of each stimulation train. Once a single unit or LFP was selected for closed-loop control it was used throughout the session.

### Electrophysiological recordings

A Multineuronal Acquisition Processor (Plexon Inc, Dallas, TX) was used to record neuronal spikes and local field potentials. Briefly, neural signals were recorded differentially, amplified (20,000–32,000X), filtered (filtering band between 400 Hz and 5 kHz), and digitized at 40 kHz. The activity of single-unit and multi-units was initially sorted using an online sorting algorithm (Plexon Inc). Local field potentials (LFPs) were acquired by band-pass filtering the raw signal (0.3–400.0 Hz), preamplified (1,000), and digitized at 1,000 Hz using a digital acquisition card (National Instruments, Austin, TX) and a multineuronal acquisition processor (Plexon).

### Behavioral assessment

Rats were placed in a rectangular open field (64.5 cm X 54 cm axes, 60 cm tall) where they were allowed to freely explore for around 40 min, and motor behaviors were recorded from a top view camera. The position of the rats in the open field was tracked using a four-colored LED placed on their head. Then positional data were extracted from digitized video recordings with custom-designed algorithms implemented in Matlab (MathWorks, USA). Instantaneous speed and instantaneous acceleration vectors were computed from the Euclidean distance between the Positional data of the rat every 1/30 seconds in the open field.

### Isolating Epochs of Locomotion and Rest

Positional data was translated into a matrix of the speed at each measured time point using the frame rate of the acquisition device. The speed data were smoothed using a moving average of ten data points. Epochs of locomotion were classified as periods of sustained speed above 1 cm/s for greater than or equal to 0.10 s. Epochs of rest were classified as periods with sustained speed below 1 cm/s for greater than or equal to 0.10 s.

### Determining significant modulation of neuronal firing rates by CLDCS

A cumulative sum algorithm with bootstrap resampling was used to detect significant changes in firing rates in neurons between the off-stimulation and CLDCS conditions. For each neuron, the firing rates were calculated over 1 second time bins, and the average binned firing rate of the off-stimulation period was subtracted from the firing rate of all time bins. Within the off-stimulation period, a 95% confidence interval around the cumulative sum of 1000 bootstrap samples of the adjusted off-stimulation firing rate was calculated. If the cumulative sum of the CLDCS condition deviated from these confidence intervals, the neuron was said to have a deviation from the previous off-stimulation activity. Within a fifteen-minute period, if five deviations in CLDCS from the off-stimulation period are detected, the neuron is identified as exhibiting a significant change in firing rate. Of these neurons, those that increased or decreased their firing rate in response to CLDCS were labeled as excitatory and inhibitory neurons, respectively.

### Principal Component Analysis of Locomotion Initiation

Bouts of locomotion initiation were identified as 3 s epochs of rest followed by 3 s epochs of movement. These bouts of locomotion initiation were then split into 0.25 s time windows, and the firing rate within these windows for all recorded neurons was calculated. Within each session, the firing rate time-series for each bout of locomotion initiation within a stimulation condition were averaged together for each recorded neuron, yielding an array of averaged time-series firing rate activity immediately before and after the initiation of locomotion for each neuron, stimulation condition, and recording session. Firing rate data were not averaged across sessions or animals. Neuronal firing rates were horizontally concatenated for each session and vertically concatenated for each stimulation condition, yielding a single matrix of T x N dimensions, with T corresponding to the number of time windows for all stimulation conditions and N to the total number of neurons monitored across all recording sessions.

Principal component analysis (PCA) was used to reduce the dimensionality of the firing rate matrix during the locomotion bouts to yield a representation of the aggregate neural state at each analyzed time point. By constructing principal components that capture maximum variance across the variables, PCA reduces and simplifies a multi-dimensional data set to extract the most meaningful information. Because each principal component is a linear combination of the original variables, which in this application are the time-series firing rate data for each neuron, the resulting scores for each principal component represent the aggregate neural activity at a given time point and stimulation condition from the recorded brain regions.

The time-series scores of the three principal components capturing maximum were selected for analysis. The selected time-series scores were plotted in three dimensions, yielding a visual representation of variance in aggregate neural activity over time. Differences in the clustering behavior in the neural activity state before and after locomotion initiation were examined using two metrics. The spread of the cluster was quantified by the volume of the minimum volume enclosing ellipsoid (MVEE) of the pre- and post-locomotion initiation neural activity states. The variance between states was quantified using the distance between the MVEE centroids for the pre- and post-locomotion initiation neural activity states.

### LFP Frequency analysis

Power spectral density (PSD) was calculated with the Welch periodogram method, using a fast Fourier transform of 512 points (frequency resolution of 1.95 Hz) and 50% overlap and a Hanning window to reduce edge effects. Beta power was defined as the average of the PSD across the beta band (10–30 Hz). The mean PSD was then computed for all the channels in four areas where the electrodes were implanted.

### Statistical analysis

The difference in locomotion parameters and beta spectral power between off-stim and CLDCS; and off-stim and pre-lesion conditions were analyzed using a two-sample t-test (Fig 2). Wilcoxon matched-pairs signed rank test was used to study differences between inhibitory firing rates, excitatory firing rates, the average volume of a fitted ellipse to the neural trajectory, and path length of neural trajectory between off-stim and CLDCS conditions (Fig 3). One-way ANOVA followed by Tukey’s Multiple Comparisons test was used to study the following differences between off-stim, OLDCS, and CLDCS conditions: locomotion parameters; beta spectral power in multiple brain areas; and distance from pre-lesion cluster centroid (Figs 4,5).

## Author Contributions

Conceptualization: APY, MALN Methodology: APY, EB Investigation: APY, EB, BR Visualization: APY, EB, BR Supervision: APY Writing—original draft: APY Writing—review & editing: APY, BR, MALN

## Data and materials availability

All data needed to evaluate the conclusions in the paper are present in the paper and/or the Supplementary Materials

